# Rawcopy: Improved copy number analysis with Affymetrix arrays

**DOI:** 10.1101/027409

**Authors:** Markus Mayrhofer, Björn Viklund, Anders Isaksson

**Affiliations:** Science for Life Laboratory, Department of Medical Sciences, Uppsala University, SE-751 85 Uppsala, Sweden

## Abstract

Rawcopy is an R package for processing of Affymetrix CytoScan HD, CytoScan 750k and SNP 6.0 microarray raw intensities (CEL files). It uses data from a large number of reference samples to produce *log ratio* for total copy number analysis and *B-allele frequency* for allele-specific copy number and heterozygosity analysis. Rawcopy achieves higher signal-to-noise ratio than commonly used free and proprietary alternatives, leading to improved identification of copy number alterations. In addition, Rawcopy visualises each microarray sample for assessment of technical quality, patient identity and genome-wide absolute copy number states.

## 1 Background

DNA copy number alteration is an important mutational process for evolution, population genomics, genetic disorders and cancer development and progression [1]. Gain and loss of gene copies may lead to extreme overexpression, absence of any functional transcript, or modest alterations in gene expression. [2] Genome-wide copy number analysis is commonly performed in hypothesis-generating genomics research, including many recent large-scale cancer studies [3]. It is also growing rapidly in clinical diagnostics as a high-resolution alternative or complement to *in situ* chromosome analysis [4, 5].

While recent advances of low-cost sequencing indicate that whole-genome sequencing may eventually become the all-in-one solution for clinical genome analysis, most copy number analysis is currently performed using microarrays [6]. Originally designed for genotyping, several brands of SNP microarrays are now marketed specifically for copy number analysis [7]. Due to their simplicity of operation and relatively manageable data analysis, the use of microarrays for copy number analysis have continued to rise in both cancer and constitutional cytogenetics [8]. It also remains standard practice in cancer genomics studies for which many thousands of samples have been published and made available for data mining by the Cancer Genome Atlas [9] and at the Gene Expression Omnibus [10].

DNA microarray signal intensities are subject to a large amount of noise and bias incurred by factors such as laboratory conditions, reagent quality, non-uniform DNA extraction efficiency along the genome and probe cross-hybridization. Systematic variation, often quantified as Median of Absolute Pairwise Differences between adjacent probes (MAPD), can be reduced using patient- or population-matched reference samples processed in an otherwise identical fashion. Systematic variation that affects samples similarly but with different strength, such as GC-content related waviness, can be further normalised for in individual samples [11].

Estimating the copy number per cell using extracted DNA from populations of cells has some important limitations. As a fixed amount of DNA is analysed rather than a fixed number of cells, any multiple of the true set of copy numbers would result in the same observation on the microarray. This is well exemplified by the aneuploidies encountered in cancer genomes, where the total amount of hybridization to the microarray does not reflect the total amount of DNA per cell in the sample. The microarray intensities are median centred to account for variation in total hybridization to the array, with the median intensity corresponding to the median copy number in the genome(s) analysed. The intensities may also be compared to those of a reference pool of samples or a patient-matched normal sample to account for systematic variation or constitutional copy number variation. Estimation of absolute copy numbers, while thoroughly explored in recent years, takes place downstream from basic normalisation of raw signal intensities and achieves estimates of the most likely absolute copy numbers given the observations [12–14].

Due to limitations in microarray sensitivity and specificity, an increase or decrease in sample DNA abundance along the genome (copy number alteration) leads to a weaker relative effect on intensity ratio, and saturation effects may differ between microarrays. The normalized intensity per probe relative to the reference is usually log transformed to equalize noise levels over different copy number states, producing the *log ratio*, a measure of DNA abundance along the genome. The relationship between log ratio and sample DNA abundance, however, differs somewhat between microarray samples and platforms [15].

Bi-allelic (SNP) probes on the microarray can, in addition to genotyping and heterozygosity mapping, be used for copy number analysis in an allele-specific manner. This results in estimates of the actual number of each parental homologous chromosomal copy per cell [12–14]. Allele-specific signal intensities per SNP probe are usually processed as estimates of the B-allele abundance relative to the total DNA abundance, called the *B-allele frequency* (BAF), ranging from near zero for homozygous A SNPs to near one for homozygous B SNPs.

Downstream analysis of log ratio and BAF includes segmentation (partitioning into segments) of the genome, for which several types of algorithms are in use. Hidden Markov Models use the expected log ratio associated with given copy number states to assign the most likely segment breakpoints given the observations, and are popular in constitutional cytogenetics where the genome can be assumed to be near-diploid and homogeneous. Circular Binary Segmentation estimates segment break points without prior assumptions of the amplitude of change incurred by copy number alterations, which is more suitable for cancer genomics where the average ploidy and purity are unknown [16].

Once segments have been defined, copy number states may be assigned to them based either on deviation from median log ratio, in which case gain and loss are defined relative to the median copy number of the genome, or using more complex analysis of segment log ratio and BAF to estimate the absolute copy numbers per cell. For Affymetrix SNP microarrays, most data analysis is performed using one of the following solutions: Chromosome Analysis Suite (or Genotyping Console for older arrays) is a proprietary solution for windows systems freely available from Affymetrix. Affymetrix Power Tools is an open source command-line alternative to Chromosome Analysis Suite, running on Linux and Mac OS. Nexus Copy Number is commercial software from Biodiscovery Inc., Hawthorne. Other free processing tools for SNP 6.0 achieve similar or lower-quality results than Affymetrix Power Tools [17]. We set out to build an open-source solution for processing of Affymetrix SNP 6.0 and CytoScan raw data (CEL files), aiming for performance better than the currently available free and proprietary alternatives and simple installation and workflow.

Rawcopy, described here, is a processing tool for Affymetrix CytoScan HD, CytoScan 750k and SNP 6.0 arrays. We demonstrate reduced systematic variation in log-ratio and BAF compared to the currently most widely used alternatives. Rawcopy is freely available as an installable R package. It is intended to provide the highest quality normalization of log ratio and B-allele frequency, suitable for downstream analysis with a range of tools. It also provides genome segmentation and several visualizations to facilitate assessment of data quality and results. The solutions presented here may also be adopted for processing of other types of microarrays and for sequencing-based copy number analysis.

## 2 Description

Rawcopy is available as an R package installable under Linux, Mac OS and Windows. Processing time per sample is 10-20 minutes depending on processor speed. The analysis may be run in parallel on multiple processor cores, with each thread requiring less than 8GB of RAM. Apart from the R package, only sample raw intensity files are required to run the analysis. Reference data are built-in and precompiled from a large number of ethnically diverse samples (Supplementary Table 1), with variations also in technical quality. The processing of new samples is described in the sections below and is schematically shown in Figure 1. B-allele frequency for SNP probes is estimated using reference sample genotypes, with normalization for total probe intensity. Total DNA abundance per probe (log ratio) is estimated by comparing total probe intensities to the reference data and normalising for sample-specific effects such as GC content and fragment length bias. Partitioning of chromosomes into segments of unchanging copy number (segmentation) is performed using the Parent-Specific CBS method [18]. Samples are then further processed to facilitate downstream analysis, including sample identity level matching, estimation of median log ratio and allelic imbalance per gene and genomic segment, and clustering and visualization of the sample set.

**Figure 1.**
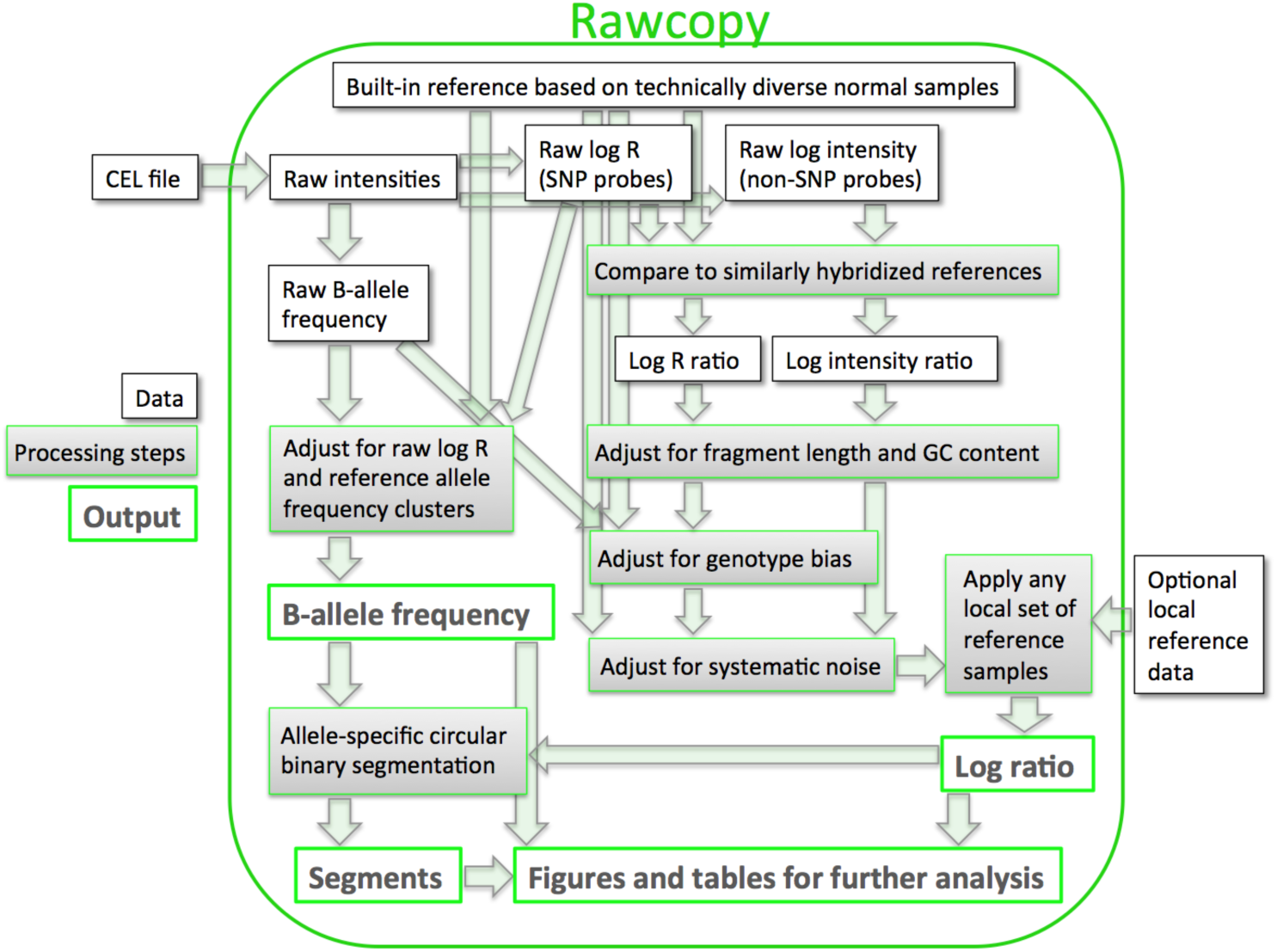
*Schematic view of Rawcopy. Grey boxes represent processing steps, white boxes represent data and green borders indicate processed data that form part of the output. The built-in reference data are used in several of the processing steps. The only required input data are the CEL files – standard format of Affymetrix microarray raw intensities. Optional input data are local reference samples and, for analysis of somatic aberrations in cancer, patient-matched normal samples. Segmentation is performed using the Parent-Specific CBS method.*

Samples are loaded into Rawcopy from raw intensity (CEL) files. Log ratio of SNP probes is based on the Euclidean sum (R) of individual allele A and B mean intensities (Ā and 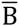, the array contains up to four physical probes for each SNP probe set and allele):

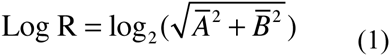

Log ratio of non-SNP probes is based on log_2_ of their raw intensities. Log ratio is composed of both SNP and non-SNP probes, these are merged during the normalization process. BAF is calculated from the raw BAF per SNP probe set:

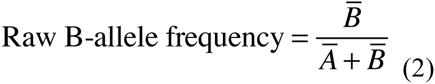

Data processed with Rawcopy are suitable for downstream analysis using a range of tools, including ABSOLUTE [14], ASCAT [12], Nexus Copy Number and TAPS [13].

### 2.1 Log ratio normalization

Log ratio of all probes is calculated using built-in reference data (listed in Supplementary Table 1). For each probe, reference log R (SNPs) or log intensity (non-SNPs) are stored in Rawcopy as a linear function of median log2 probe intensity (the amount of hybridization to the microarray) and the raw experimental variation (raw MAPD) of the reference samples, as shown in Figure 2A. After subtracting the reference value for each probe, given the median hybridization and MAPD of the new sample, log ratio represents logarithmized observations of hybridization relative to the reference level.

**Figure 2.**
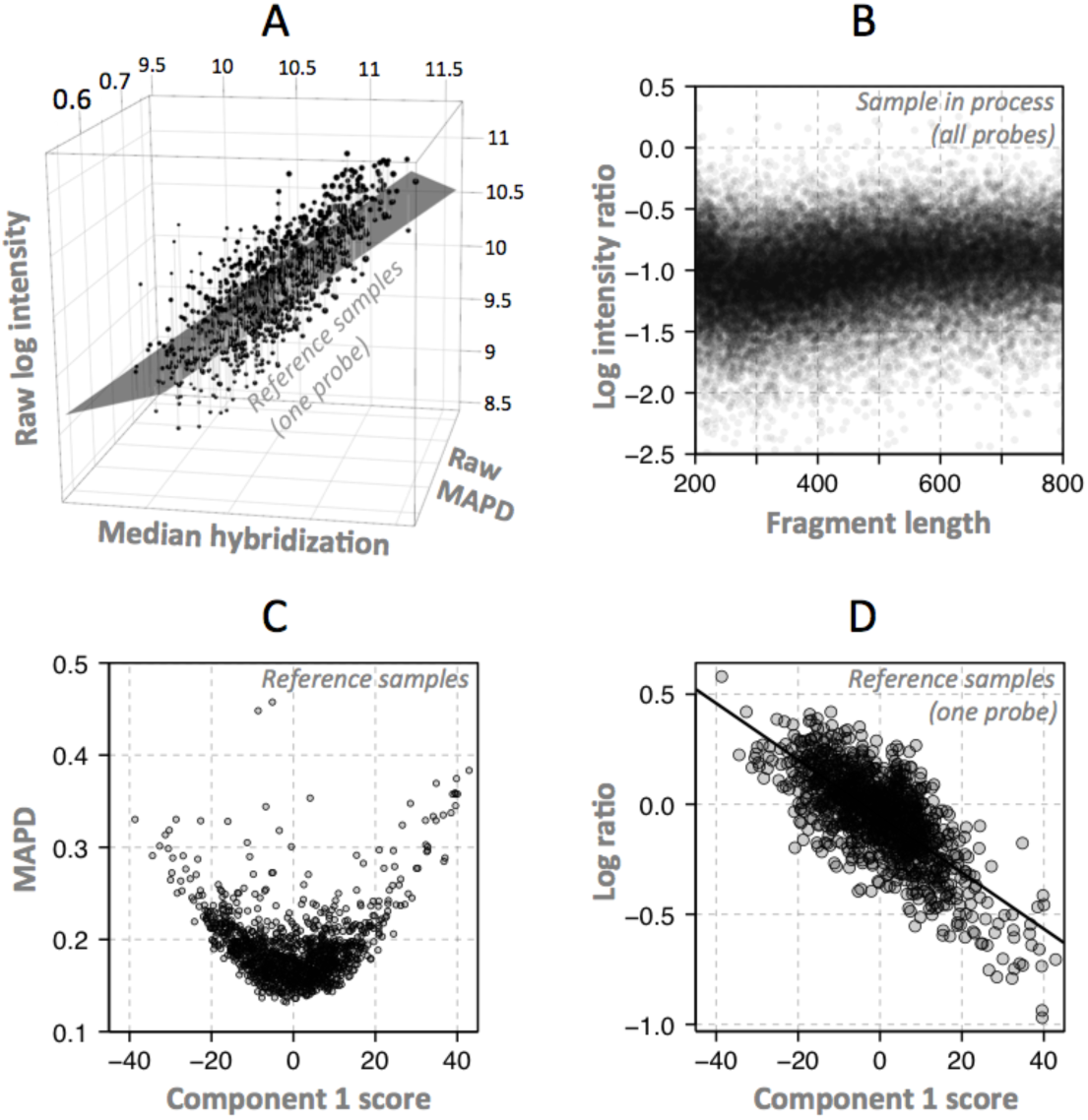
*Log-ratio processing.* ***A**: For each probe, the expected raw intensity is modelled as a linear function of the median hybridization level and raw median absolute pairwise difference (raw MAPD) of the reference samples. Robust linear regression is used to minimize influence of copy number variation, which is assumed to be present in relatively few reference samples for each probe. Only female reference samples were used for the X chromosome and only male samples for the Y chromosome. When processing a new sample, log ratio of each marker is calculated from the difference between the new observation and the reference function value given the median hybridization and raw MAPD of the new sample.* ***B**: Example of fragment length bias after the previously described step. Fragment length bias and GC content bias may differ between samples and are therefore only partially corrected for using the reference sample data. Remaining fragment length bias and GC content bias (which may be nonlinear) are corrected for simultaneously by equalizing the median log intensity ratio within percentiles of fragment length and GC content.* ***C**: After subjecting reference sample log ratio to multidimensional scaling (MDS), the MDS component scores of reference samples indicated systematic differences between samples, as samples which deviated from the mean along any component were associated with higher MAPD (noise).* ***D**: Example of a probe where log ratio correlates with the first MDS component score of each sample.*

Fragment length and GC content bias, which differs between samples (Figure 2B), are then adjusted for separately in each sample by median-centring the log-ratio within percentiles of both fragment length and GC content. Raw BAF is also used to adjust the log ratio of SNPs linearly for correlation between genotype and log ratio in the reference data.

To describe additional systematic variation in the reference material, autosomal log ratio of all reference samples were subjected to multidimensional scaling (MDS), compressing them from one dimension per probe into a few components. The vast majority of variation in the reference material was expected to be noise rather than copy number alterations, and this was also indicated by the data as samples that deviated from the average along any component were associated with more noise (higher MAPD, Figure 2C). For most probes, the log ratio of reference samples correlated with their component scores (Figure 2D). This correlation was weaker for each additional component in the MDS (data not shown). Six components were selected as a balance between reducing noise and minimizing data storage in Rawcopy. Linear functions of these six components (for which each sample has a score) are stored in Rawcopy and used to reduce noise for each probe by subtracting the function value, given sample component scores, from the observed log ratio. When processing a new sample, the score giving the lowest MAPD is determined and used for each of the six components.

If a local set of reference samples is available, it can be used to further reduce noise and waviness, which may partially be specific to reagents, cell types and other laboratory conditions. This is done by subtracting the median local reference log ratio from that of the current sample, for each probe.

### 2.2 B-allele frequency estimation

Due to background hybridization and the possibility of unequal specific and non-specific hybridization of the two alleles, the raw BAF defined in Formula 2 cannot be assumed to accurately represent the true BAF. In Rawcopy, the raw BAF associated with each normal diploid genotype (AA, AB and BB) in the reference material are stored in Rawcopy as functions of the “log R” defined in Formula 1. Examples of SNPs with well-defined genotype clusters are shown in Figure 3A-B.

**Figure 3.**
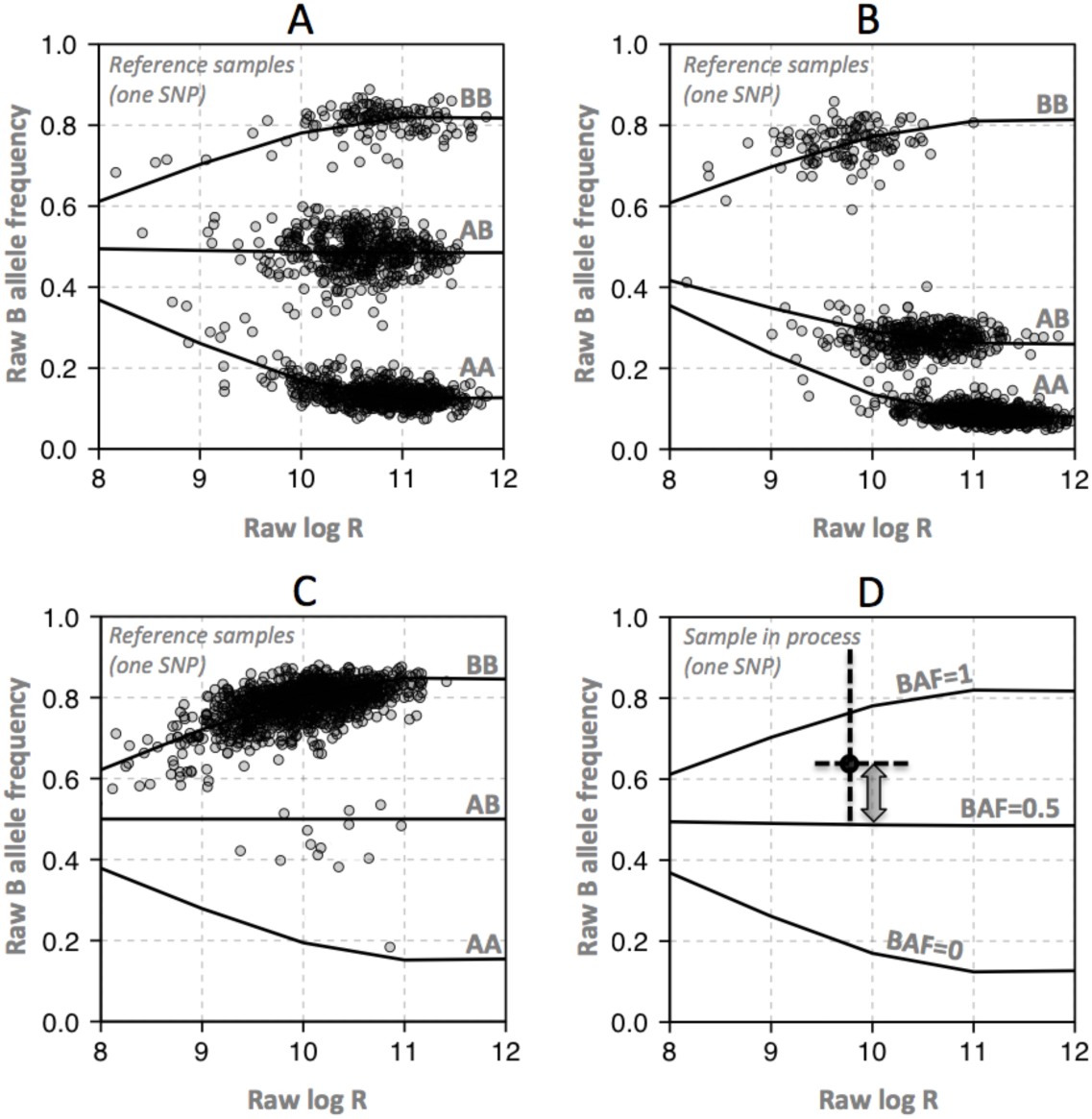
*BAF estimation.* ***A**: Reference samples were used to express raw B-allele frequency as functions of raw log R given genotypes AA, AB and BB, for all SNPs. To reduce extrapolation problems, a characteristic function was designed based on the general behaviour of the data, with parameters that could be fitted for individual SNPs. SNPs with good separation of clusters and with each cluster populated by multiple samples were considered high-quality.* ***B**: Robustness against hybridization bias between genotypes was confirmed by visually inspecting a large number of SNPs.* ***C**: SNPs with poor separation of clusters and/or difficulty of defining cluster centres were considered lower-quality reference data. For them, the mirror image around 0.5 of the most well-defined homozygous cluster was used instead of the poorly represented homozygous genotype, and a straight line at 0.5 was used for heterozygous SNPs.* ***D**: When processing new samples, BAF is calculated using the stored function parameters for each SNP and genotype cluster, and the raw BAF and log R of the new sample. In this example, raw BAF (black dot) appears above the function line for the AB genotype. BAF is therefore calculated linearly from the observation relative to the AB (BAF=0.5) and BB (BAF=1) function lines, yielding a value near 0.75.*

A subset of SNPs (35% of CytoScan, 44% of SNP 6.0) displayed either poorly separated clusters or no variation in genotype among the reference samples as a result of too low minor allele frequency. The reference data for those SNPs are therefore considered lower quality. An example of such a SNP is shown in Figure 3C. Inclusion of lower-quality SNP data is optional in Rawcopy. Heterozygosity rates for SNPs associated with high- and low-quality reference data are presented in Supplementary Figure 1. When processing new samples, Rawcopy estimates the BAF of each SNP as shown in Figure 3D, given the observed “log R” and raw BAF.

### 2.3 Genome segmentation

The segmentation step available in Rawcopy uses the PSCBS package [18]. Once segment break points have been determined, segments are annotated with median log ratio, number of probes, genes and cytoband. Allelic imbalance for genomic segments is quantified in the same way as in TAPS [13]. Segment tables are written to tab-separated text files for browsing and further analysis.

### 2.4 Data visualization

After processing of log ratio, BAF and segmentation, whole-genome and chromosome-wise figures are plotted for each sample. These allow the user to assess technical quality of individual samples such as total hybridization level and quality of the physical array, and get an extensive overview of chromosomal alterations as shown in Figure 4.

**Figure 4.**
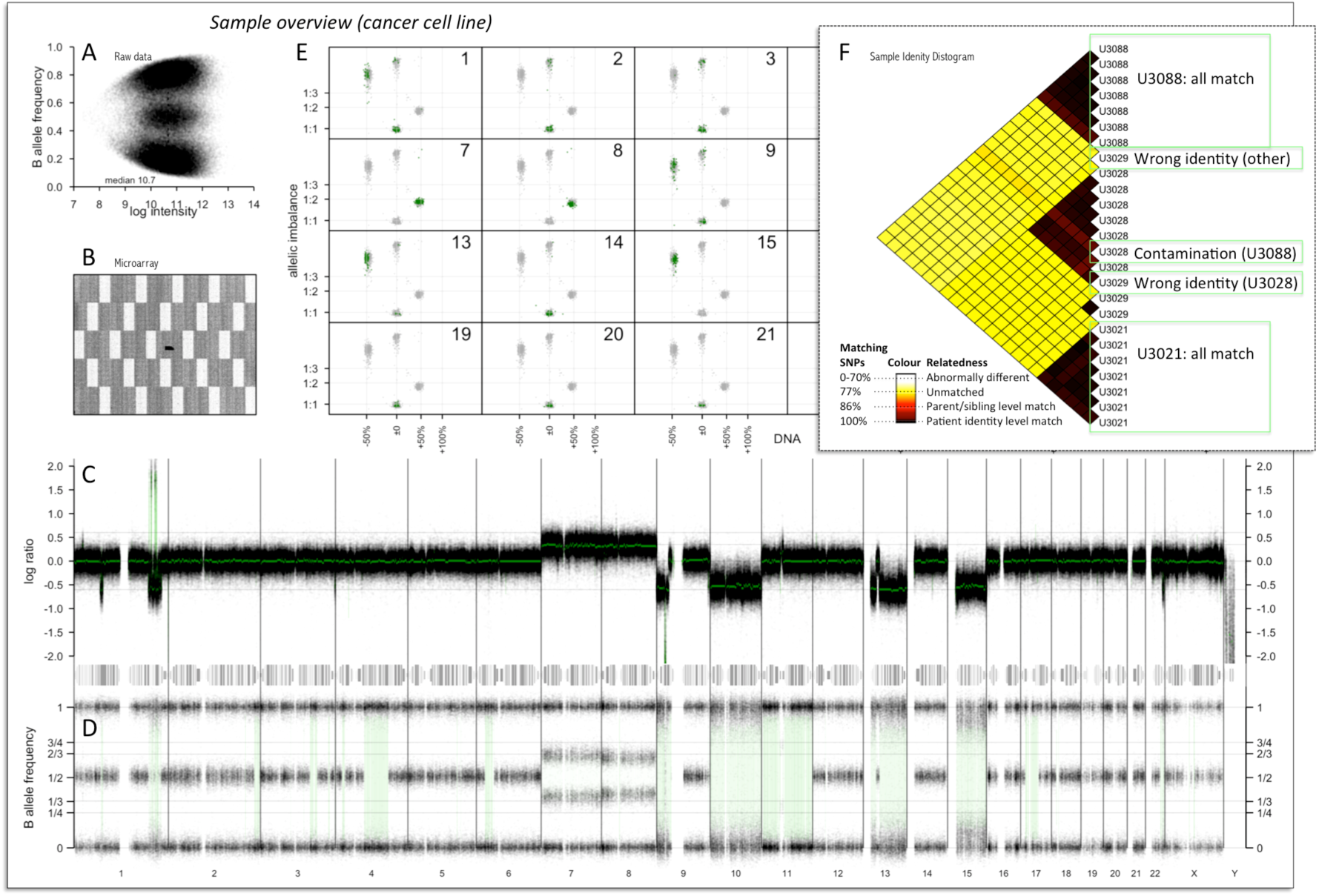
*Rawcopy visualization per sample (A-E) and data set (F)* ***A**: A scatter plot of raw log intensity versus raw B allele frequency indicates the overall hybridization and genotype cluster distribution of the sample.* ***B**: Raw signal intensity throughout the microarray may indicate uneven hybridization and array quality issues.* ***C**: Log ratio indicates alterations in total copy number along the genome. Segments are marked in green and small amplifications and deletions are highlighted for visibility.* ***D**: B-allele frequency indicates the genotype of each SNP probe. Most copy number alterations lead to allelic imbalance indicated by absence of the middle band of AB genotype SNPs. Allelic imbalance and homozygous segments are highlightedfor visibility.* ***E**: Scatter plots of segmented DNA abundance estimates and allelic imbalance, with individual chromosomes highlighted relative to the rest of the genome, indicate absolute copy number per cell and any heterogeneity in the cell population.* ***F**: Genotype data from all samples in the batch are cross-matched and patient identity level dissimilarity scores are computed. This example shows multiple cancer cell lines from four tumours. Two examples of erroneous sample identity are indicated by the distogram; one U3029 sample matches U3028 instead and one U3029 sample matches none of the other samples. In addition one U3028 sample shows some cell or DNA contamination from U3088.*

To visualize the copy number throughout the genome, scatter plots of total copy number and allelic imbalance are shown throughout each chromosome relative to the rest of the genome (Figure 4E). This is equivalent to a previously published solution [13, 15] and can be used to indicate the absolute number of copies involved in copy number alterations, mosaicism, and the ploidy and purity of cancer samples. In the scatter plots, the median log ratio of each segment (about 1MB long) is transformed into estimates of DNA abundance relative to the median of the current sample. Cancer cell line samples [19] were used to set the expected log ratio given 50% loss (log ratio: -0.6), 50% gain (log ratio: 0.35) and 100% gain (log ratio: 0.6) of DNA abundance relative to the median of the genome. (Individual samples may deviate slightly from this model for technical reasons.) Allelic imbalance is measured for each segment by first clustering abs(BAF-0.5) on two means, representing the separation of heterozygous and homozygous SNPs (BAF_het_ and BAF_hom_), then quantifying the separation of BAF_het_ relative to BAF_hom_:

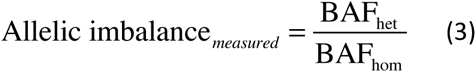

Assuming heterozygous SNPs exist and are separated into two bands due to imbalanced copy number, allelic imbalance represents estimates of the difference in homologous copies (H) relative to the total copy number:

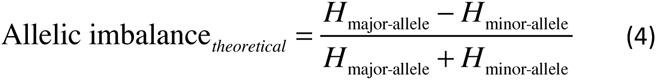

Due to the systematic variation in BAF, segments where the copy number is balanced or homozygous result in measured allelic imbalances just above 0 or below 1, respectively (Figure 4E). The scatter plots are annotated with the expected allelic imbalance given a 1:1, 1:2 and 1:3 ratio of homologous copies.

The pairwise genotype dissimilarities of all samples processed together are plotted as shown in Figure 4F. BAF is discretized into B-allele presence (1 if BAF≥0.2, else 0), reducing the effect of systematic and copy number variation while largely retaining genotype information on sample identity level. Pairwise dissimilarities (sum of differences in BAP) between samples are then visualized in a distogram [20], using a colour gradient based on observed dissimilarities between related and unrelated members of HapMap CEU. In addition to validating sample identities relative to one another, the sample identity distogram may indicate cell or DNA contamination.

## 3 Performance

Two large sets of publicly available samples were acquired from the Gene Expression Omnibus (GEO) for systematic benchmarking of performance relative to some of the most commonly used free and proprietary processing tools. For the Affymetrix SNP 6.0 platform, a set of 947 cancer cell lines [21] published by the Broad Institute, Massachusetts (GEO accession number: GSE36138) was analysed using Affymetrix Power Tools, Nexus Copy Number and Rawcopy. For the Affymetrix CytoScan HD platform, a set of 231 hepatocellular carcinomas [22] published by the Gachon University of Incheon, South Korea (GEO accession number: GSE54504) was analysed using Affymetrix Power Tools, Nexus Copy Number, Chromosome Analysis Suite and Rawcopy. In addition, this set of samples was analysed with Rawcopy using the included matched normal samples as local reference data to further reduce noise.

### 3.1 Reduced log ratio noise relative to true signal

The most commonly used metric of technical quality in microarray copy number analysis is the MAPD which estimates the amplitude of log ratio noise in a way that is largely unaffected by copy number alterations. As the majority of adjacent measurements of log ratio should be the same, the median of their absolute pairwise differences are frequently relied upon to compare technical quality across samples with different distributions of copy number states. However as some methods employ normalization steps that alter the distribution of the data, such as quantile normalization, the MAPD may not be comparable across different processing tools. In cancer samples with large copy number alterations, MAPD may be adjusted based on the observed effects of copy number alteration on log ratio. We defined the signal-adjusted pairwise difference (SAPD) as the MAPD divided by the effect ∆ of copy number alteration on log ratio with the current processing tool, relative to the average effect 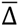 over a set of different processing tools (given the same sample and copy number alteration):

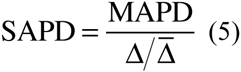

MAPD and SAPD were calculated for each sample and each processing tool in the evaluation. For each sample in the evaluation data, the two autosomes with the highest and lowest median log ratio was selected for calculating ∆ with all processing tools. Samples with little or no evidence of copy number alteration were removed (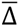 less than 0.2). SAPD of the evaluation samples are shown for Rawcopy and the most commonly used current processing tools in Figure 5.

**Figure 5:**
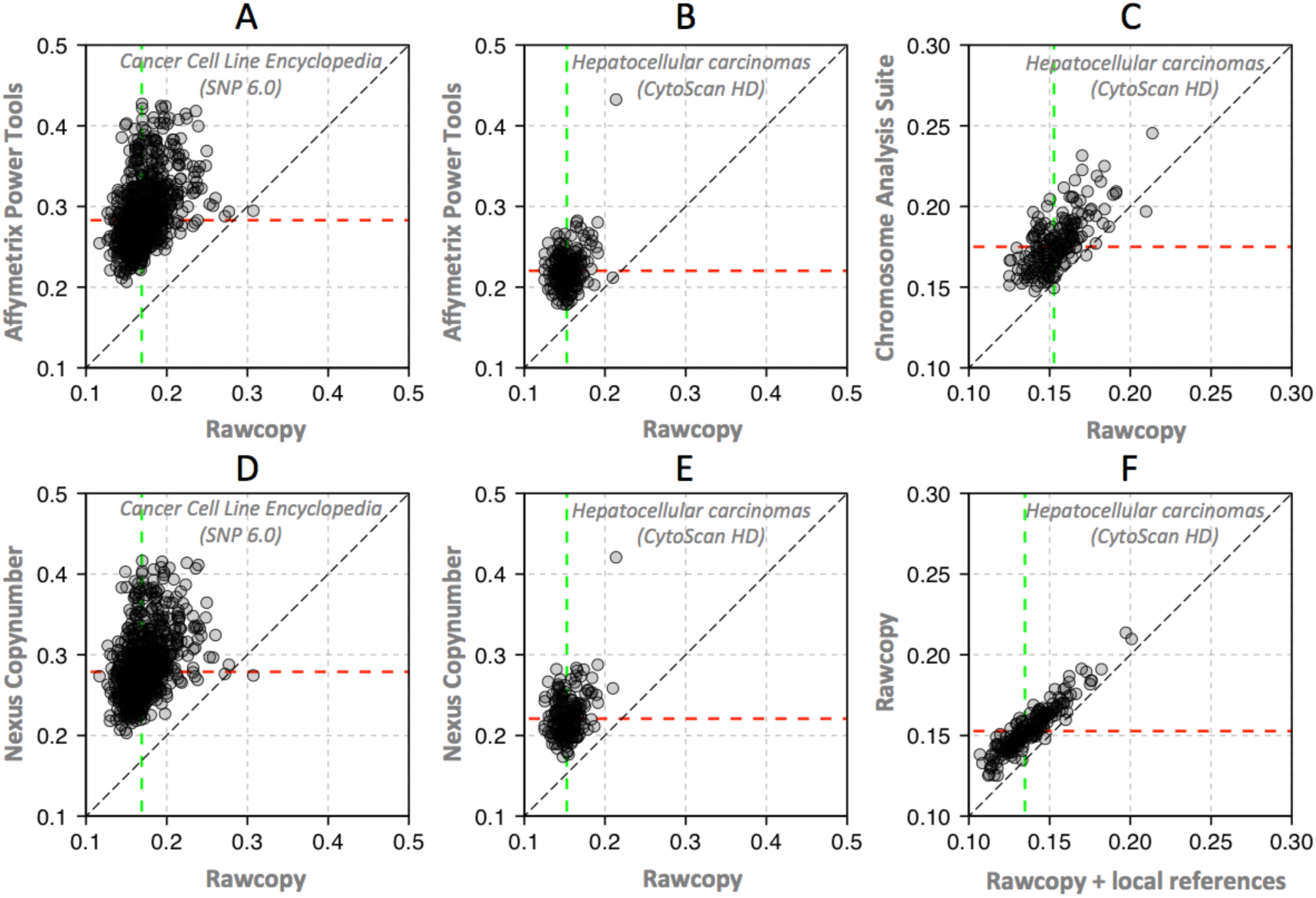
*Noise estimates for Rawcopy and other processing tools.* *Signal-adjusted Pairwise Difference (SAPD) of evaluation samples with Rawcopy and commonly used alternatives (lower values are better). Dashed lines indicate median SAPD in the evaluation set of samples. Rawcopy achieved significantly better SAPD than each of the alternatives for both the CytoScan HD and SNP 6.0 evaluation sets (p<2.2*10^−16^, paired Student’s T-tests). Rawcopy median SAPD was 0.15 for CytoScanHD and 0.17 for SNP 6.0.* ***A**: Affymetrix Power Tools and SNP 6.0. Median SAPD: 0.28.* ***B**: Affymetrix Power Tools and CytoScan HD. Median SAPD: 0.22.* ***C**: Chromosome Analysis Suite and CytoScan HD. Median SAPD: 0.18.* ***D**: Nexus Copy Number and SNP 6.0. Median SAPD: 0.28.* ***E**: Nexus Copy Number and CytoScan HD. Median SAPD: 0.22.* ***F**: Use of local reference samples (patient-matched normals used as a local reference data set) further reduced Rawcopy SAPD. Median SAPD: 0.13.*

### 3.2 Improved estimates of allele frequency

Rawcopy and Nexus Copy Number both produce the BAF (estimates the relative B allele frequency in the DNA for each SNP), but Nexus Copy Number truncates the data at zero and one. Affymetrix processing tools produce Allele Difference (sometimes called Allele Peaks) instead, representing the difference between log_2_(B) and log_2_(A). These different approaches result in similarly useful but somewhat different looking allelic data, shown for a representative evaluation sample in Figure 6. Rawcopy and Nexus achieve better separation and stability of SNPs with near-equal abundance of the A and B allele compared to Chromosome Analysis Suite (6A). Rawcopy also achieves the best separation of homozygous and near-homozygous SNPs (6B-C). Rawcopy BAF is less skewed by total DNA abundance than Nexus BAF (6B,D).

**Figure 6:**
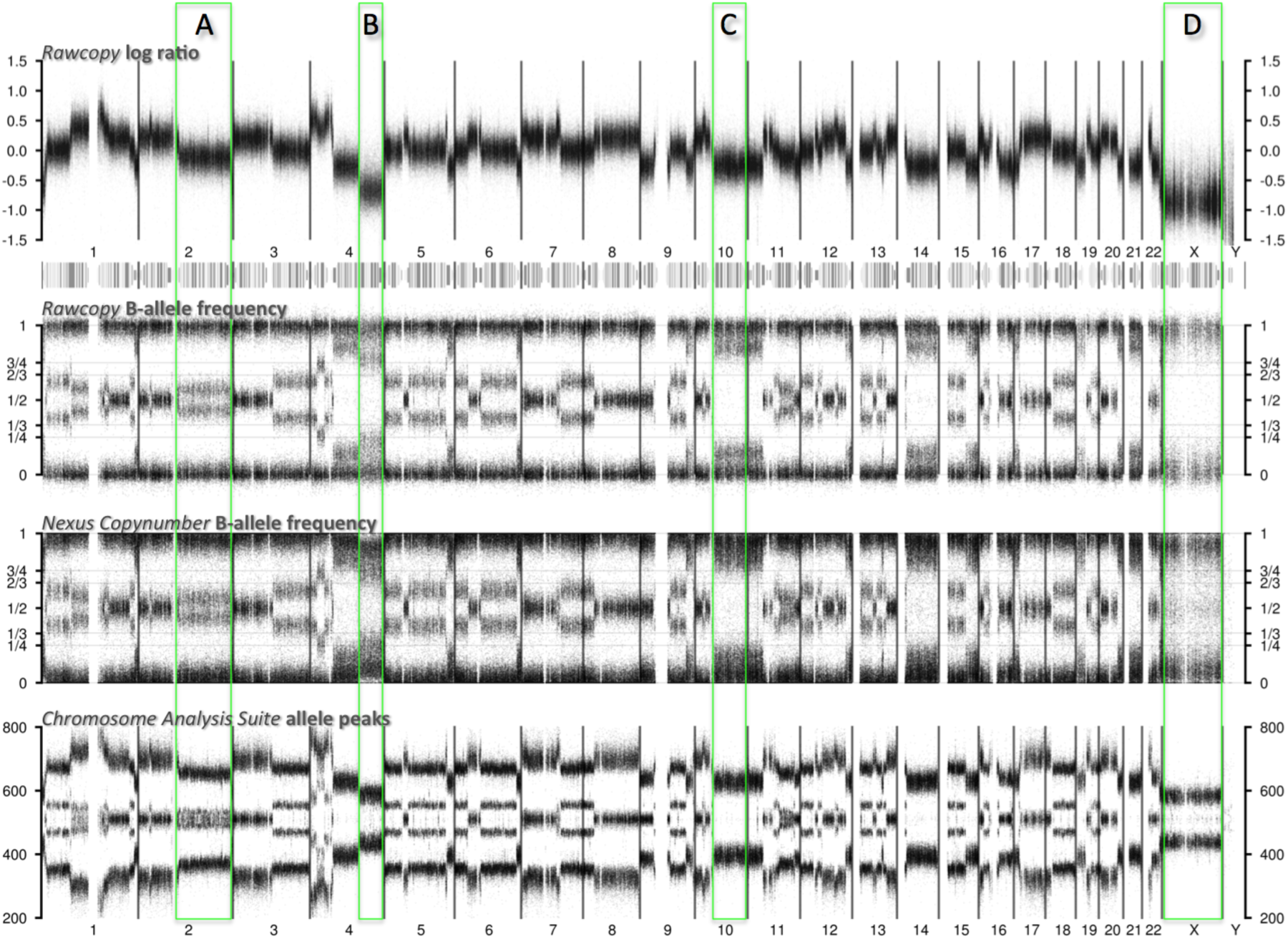
*Representative example of log ratio, BAF and Allele Peaks from Rawcopy, Nexus Copy Number and Chromosome Analysis Suite.* ***A**: In the BAF tracks the middle band of heterozygous SNPs has split, indicating some allelic imbalance. Chromosome Analysis Suite allele peaks do not show this clearly throughout the chromosome arm, oscillating instead between a balanced (one middle band of SNPs) and unbalanced profile.* ***B**: At low total copy number, Nexus BAF shows some bias as homozygous SNPs migrate towards the middle. With Rawcopy, homozygous SNPs remain steadily near 0 and 1, indicating that such bias is successfully normalized for. Chromosome Analysis Suite allele peaks do not clearly indicate separation between homozygous (outer bands) and heterozygous (inner bands) SNPs.* ***C**: Rawcopy BAF shows distinction between homozygous and heterozygous SNPs, indicating that the abundance of the minor allele is relatively low but not zero.* ***D**: With Rawcopy, all SNPs of the homozygous X chromosome (of this male sample) appear near 0 or 1. With Nexus, BAF appears closer towards the centre as the limited amount of DNA leads to less separation of A and B clusters than expected given normal copy number. In addition, with Rawcopy, SNPs with low hybridization are less prone to appear heterozygous than with Nexus or Chromosome Analysis Suite (where seemingly heterozygous SNPs appear despite a homozygous X chromosome).*

### 3.3 Significance

Rawcopy makes copy number analysis easy to set up, as only installation of the R-package is required to start processing CEL files. The large built-in reference data lead to better quality of the copy number data, i.e. reduced noise relative to signal and more accurate BAF, compared to the most widely used free and proprietary alternatives. A noise level threshold (MAPD or similar) is the most commonly used quality metric for SNP microarrays, but differences in the signal distribution achieved by different processing tools preclude direct comparison of MAPD between them. MAPD may be corrected for such differences, allowing systematic evaluation of noise between processing tools. Such correction (i.e. use of SAPD over MAPD) is not suggested when processing new data as it is intended for comparing tools, not samples. All new samples cannot be expected to harbour copy number alterations of sufficient length and amplitude to make signal-to-noise assessment practical. Our evaluation does indicate that if a certain signal-to-noise ratio is required, samples have a better chance of passing such requirements with Rawcopy compared to the current alternatives. In addition, as less noise leads to more reliable data per probe, the effective genomic resolution is improved for most samples as fewer probes may be required to observe statistically significant alterations in DNA abundance.

Figures generated by Rawcopy allow immediate assessment of individual sample quality and copy number profile. Technical issues such as DNA quality and microarray fabrication errors can be identified and recognized on individual arrays and the sample identity distogram can identify mislabelled samples and reveal DNA or cell contamination. Rawcopy is suitable for copy number and heterozygosity analysis of both tumour and constitutional DNA.

Absolute allele-specific copy numbers may be estimated using scatter plots of median log ratio versus allelic imbalance for genomic segments, even for cancer samples with extensive aneuploidy, low purity or subclonal heterogeneity [13]. Copy number analysis of DNA extractions from populations of cells is ambiguous in nature as there may be more than one set of absolute copy numbers per cell that would explain the composition of DNA. With Rawcopy, we have built a tool capable of revealing rich information about the copy number profile, but without applying any automatic classification or interpretation (such as purity and ploidy for cancer samples) that could fail for samples with an unforeseen chromosomal setup. Arguably such interpretations should be done taking into account the specific disease and its frequency of specific karyotypes and genome doublings, as done in ABSOLUTE [14], but even then many samples cannot be correctly resolved as alternative solutions may be near-equally plausible. If the interpretation impacts diagnosis or other clinical decisions, cell-based chromosome or ploidy analysis may be motivated as a complement to the microarray. If specific estimates of genome-wide of absolute allele-specific copy numbers (i.e. numeric data for further analysis) are required, the output generated by Rawcopy is suitable for downstream analysis tools such as ABSOLUTE, ASCAT or TAPS. The reduction of systematic copy number-induced BAF bias achieved in Rawcopy also makes BAF a more accurate representation of true allele ratio and more likely to fit theoretical models of cell fractions with certain copy number states.

**List of abbreviations**
BAF: B allele frequency
CBS: Circular binary segmentation
GEO: Gene Expression Omnibus
HMM: Hidden Markov model
MAPD: Median absolute pairwise difference
MDS: Multidimensional scaling
PSCBS: Parent-specific CBS
SAPD: Signal-adjusted pairwise difference

## Availability

Rawcopy is free software and may be redistributed and/or modified under the terms of the GNU General Public License as published by the Free Software Foundation; version 2. Installation, execution and access are described at http://rawcopy.org.

## Authors’ contributions

MM and BV designed and developed Rawcopy. MM and AI wrote the manuscript. AI supervised the project. The final manuscript was read and approved by all authors.

## Description of additional data files

Additional File 1: Supplementary Table 1 and Supplementary Figure 1

## Acknowledgements

The authors acknowledge funding from Uppsala County Council, the Swedish Cancer Research Fund and Lions Cancer Research Fund Uppsala-Örebro. Ann-Charlotte Thuresson and the Department of immunology, genetics and pathology, Uppsala University, are acknowledged for their contribution of CytoScan HD reference samples. Affymetrix provided additional HapMap CytoScan HD and CytoScan 750k reference samples. The Cancer Genome Atlas project is acknowledged for non-cancer SNP 6.0 reference samples.

